# Unsupervised embedding of single-cell Hi-C data

**DOI:** 10.1101/257048

**Authors:** Jie Liu, Galip Gürkan Yardımcı, Dejun Lin, William Stafford Noble

**Affiliations:** Department of Genome Sciences, University of Washington; Paul G. Allen School of Computer Science and Engineering, University of Washington

## Abstract

Single-cell Hi-C (scHi-C) data promises to enable scientists to interrogate the 3D architecture of DNA in the nucleus of the cell, studying how this structure varies stochastically or along developmental or cell cycle axes. However, Hi-C data analysis requires methods that take into account the unique characteristics of this type of data. In this work, we explore whether methods that have been developed previously for the analysis of bulk Hi-C data can be applied to scHi-C data. In this work, we apply methods designed for analysis of bulk Hi-C data to scHi-C data in conjunction with unsupervised embedding. We find that one of these methods, HiCRep, when used in conjunction with multidimensional scaling (MDS), strongly outperforms three other methods, including a technique that has been used previously for scHi-C analysis. We also provide evidence that the HiCRep/MDS method is robust to extremely low per-cell sequencing depth, that this robustness is improved even further when high-coverage and low-coverage cells are projected together, and that the method can be used to jointly embed cells from multiple published datasets.

## 1 Introduction

High-throughput DNA sequencing technology now allows us to reliably measure many genomic features at the single cell level, including RNA-seq for RNA expression [1], ATAC-seq for chromatin accessibility [2], and Hi-C for 3D genome architecture [3]. In principle, these technologies provide scientists with the opportunity to understand many aspects of fundamental functional processes in the cell, including gene regulation and DNA replication. However, such understanding likely cannot be achieved via analytical methods that fail to accurately capture the complexities of these types of data.

In particular, single-cell assays introduce a new axis of variation—cell-to-cell variability—that is not directly observable in data derived from a bulk sequencing assay that profiles an aggregate of many cells. In some cases, independent labeling of cells via, e.g., fluorescence activated cell sorting (FACS) can identify cellular states for use in a supervised analysis. This type of labeling experiment, however, is more costly, lower throughput, and only applicable in a limited range of experimental systems. Thus, in most experimental settings, automated methods for characterizing cell-to-cell differences are useful.

A variety of methods have been developed for the unsupervised analysis of single-cell RNA-seq data. These methods employ a wide variety of analytical techniques. Monocle uses independent component analysis, followed by a minimum spanning tree to recover lineages [4]. Its successor, Monocle 2, uses a reversed graph embedding algorithm [5]. Other methods for scRNA-seq analysis use principal components analysis and hierarchical clustering (pcaReduce) [6], clustering based on approximate nearest neighbors (scmap) [7], biclustering (BackSpin) [8], a latent variable model (svLVM) [9], bifurcation analysis of a nearest neighbor graph (Wishbone) [10], a hidden Markov model coupled with probabilistic Kalman filtering (TASIC) [11], and a latent Dirichlet allocation model (cellTree) [12].

In this work, we focus on unsupervised methods for characterizing cell-to-cell variability in single-cell Hi-C (scHi-C) data. We choose to focus on Hi-C data because of the relative sparsity of existing methods for analyzing this type of data, and because we believe that scHi-C data will become increasingly valuable for the study of diverse developmental and disease-related processes. None of the methods developed for analysis of scRNA-seq data can be applied directly to scHi-C data.

For scHi-C data, most existing analyses of cellular heterogeneity have focused on the so-called “contact distance profile” of each individual cell (defined below). The output of a Hi-C experiment is typically summarized in a Hi-C matrix M, where rows and columns of *M* correspond to fixed-width genomic loci (typically using bin sizes of 40 kb or 100 kb). In this matrix, the value *M*_*i, j*_ is an integer count (or a normalized version thereof) representing the number of observed paired-end reads uniquely linking locus *i* to locus *j*. We refer to these paired-end reads as “contacts” and to matrix *M* as a “contact matrix.” With this input, the contact probability *P(s)* is defined as the proportion of intra-chromosomal contacts that link pairs of loci separated by s bins along the genomic axis:

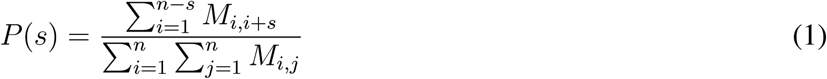

Working with bulk Hi-C, Naumova et al. showed that the contact probability function differs between mitotic and interphase cells [13]. In the single-cell setting, the data consists of a series of matrices, so that *M_i,j,k_* is the contact count for loci *i* and *j* in cell *k*. In this setting, we can compute a contact probability function separately for each cell. Accordingly, Nagano et al. used the values of *P(s)* for *s* = 1, …, *n* as a vector representation of individual cells in a scHi-C experiment. They defined the proportion of near contacts *p*near = 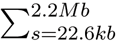 *P(s)* and the proportion of mitotic contacts *p*mitotic = 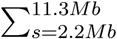 *P(s)*, and then separated the single cells into different cell-cycle stages by thresholding pnear and pmitotic. Nagano et al. demonstrated that the resulting cell-cycle phases largely agree with labels derived from FACS labeling [14]. The contact probability profile approach has also been adopted by other scHi-C analyses, including [15] and in the analysis of data generated by an alternative scHi-C protocol [16].

The current work is motivated in part by the observation that the contact probability profile captures only one aspect of genome 3D architecture. In particular, this profile focuses on the DNA self-interaction profile and omits structural information, such as where individual contacts lie along the genome, as well as higher-order properties of DNA structure such as loops and domains [17, 18]. We are interested in exploring alternative analytical techniques for scHi-C data that retain a richer representation of the underlying data.

Although scHi-C analyses have relied almost exclusively on the contact probability profile, a variety of methods have been developed for evaluating the reproducibility of bulk Hi-C data sets (reviewed in [19]). The first such method, HiCRep, first smooths the Hi-C contact matrix and then computes a weighted similarity measure separately at each genomic distance [20]. GenomeDISCO treats the Hi-C matrix as a weighted network, applying a random walk to smooth the matrix and then computing an L1 similarity score [21]. HiC-Spector transforms the Hi-C contact map to a Laplacian matrix and then summarizes the Laplacian by matrix decomposition [22]. Finally, QuASAR calculates an interaction correlation matrix, weighted by interaction enrichment [23]. Critically, each of these four reproducibilty measures was specifically designed to capture the biologically relevant signal in a noisy Hi-C data set, while reducing “uninteresting” similarities. Accordingly, we hypothesized that these methods might be useful for characterizing cell-to-cell variability in scHi-C data, and we set out to test this hypothesis empirically. Our tests focus on HiCRep, GenomeDISCO and HiC-Spector, because we failed to get the QuASAR software to run on our data sets.

Our experiments suggest that HiCRep performs significantly better than GenomeDISCO and HiC-Spector when the methods are used in conjunction with multidimensional scaling (MDS) to embed scHi-C data into two dimensions. In addition, the HiCRep+MDS method approach outperforms contact-distance-based methods for separating cells at different stages of the cell cycle. The HiCRep-based embedding approach is also quite robust with respect to the number of contacts required per cell: even with as few as 10,000 contacts per cell, the method accurately orders cells by their cell cycle phase. We are particularly interested in characterizing cells with even fewer counts, because the newer scHiC assay based on combinatorial barcoding produces data from many single cells but with lower average sequencing depth [16]. Accordingly, we demonstrated that even with very shallow sequencing (1k contacts per cell), the cell cycle can be called accurately as long as the cells are projected jointly with other, more deeply sequenced cells. Finally, we demonstrate that our proposed embedding approach supports joint projection of multiple scHi-C data, offering the potential to compare cells across different studies that capture diverse cell types or developmental stages.

## 2 Methods

### 2.1 Data sets

Two sets of single cell Hi-C data sets from recent studies are used in our experiments. Both data sets are from the mouse cells and were mapped to the mouse genome assembly mm9.

#### Cell-cycle dataset

The first set of scHi-C data [14] consists of 1171 scHi-C contact maps from F1 hybrid 129 × Castaneus mouse embryonic stem cells (ESCs). These cells were grown in 2i medium without feeder cells, tested for mycoplasma contamination, and screened based on Oct-3/4-immunoreactivity, so that there is no differentiation among the cell population. The cell-cycle phase of each cell was determined based on levels of the DNA replication marker geminin and DNA content measured via FACS. This analysis assigned 280 cells to the G1 phase, 303 cells to early-S, 262 cells to mid-S, and 326 cells to late-S/G2. The scHi-C libraries were sequenced to produce 0.89 million reads per cell on average, with per-cell coverage ranging from a minimum of 0.63 M to a maximum of 1.05 M. For each cell, uniquely mapping read pairs were aggregated into contact matrices with bins of 500 kb. In the resulting matrices, the total number of distinct contacts per cell ranges from 20–654 k with a median 273 k.

#### Oocyte-zygote dataset

The second set of scHi-C data contains 40 transcriptionally active immature oocytes (nonsurrounded nucleolus, NSN), 76 transcriptionally inactive mature oocytes (surrounded nucleolus, NSN), 30 maternal nuclei from zygotes and 24 paternal nuclei from zygotes. Both the maternal and paternal nuclei from zygotes are predominantly in the G1 phase. The number of contacts from the four types of cells are, respectively in the ranges of [1.4 k, 1.65 M], [1.2 k, 1.03 M], [4.8 k, 288 k], and [2.9 k, 294 k] with medians 66 k, 235 k, 97 k, and 117 k, respectively. Note that the scHi-C protocol used to generate this dataset differs markedly from the one used for the cell-cycle dataset, resulting in approximately ten-fold more contacts per cell.

### 2.2 Similarity and distance measures for scHi-C contact maps

In this study, we consider one distance measure and three similarity measures for scHi-C contact maps.

The distance is based on the contact distance profile of the Hi-C contact maps, described by Equation 1. To compute the distance, we first build a vector representation *C* of the contact distance profile (CDP) for each chromosome *c* of each cell *k*, where one CDP entry corresponds to one contact distance, i.e.,

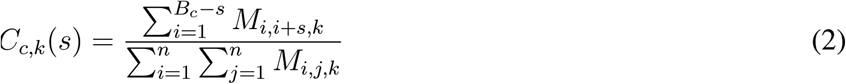

where *s* is the distance in units of the contact matrix bin size (i.e., 500 kb in this work), and *B*_*c*_ is the number of bins in the chromosome. We sum *C*_*l, k*_(*s*) over all chromosomes and then normalize to get the CPD for each cell:

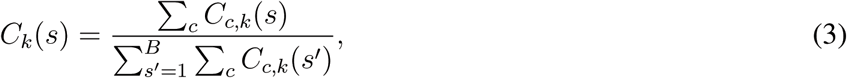

where *B* is the number of bins in the largest chromosome. For shorter chromosomes, the contact profile values for bins beyond the end of the chromosome are set to zero. Finally, we compute the distance between two cells using the Jensen-Shannon divergence (JSD) between the CDPs:

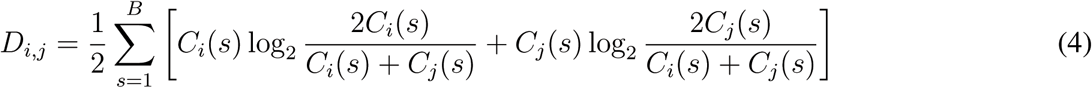

We term this contact-distance-based measure “CDP-JSD.”

The other three measures directly estimate the similarity from the Hi-C contact matrices. All three methods have been described previously, so we only provide a brief overview here.

HiCRep [20] consists of three steps. First, the Hi-C matrix is smoothed using the 2D equivalent of a sliding window smoothing operation: the observed contact count for loci *i* and j is replaced by the sum of contacts between loci in fixed-size windows around *i* and *j*. In this work, we follow the recommendations from the original HiCRep paper and use a window size of 3 (i.e., *i* − 1, *i, i* + 1) for Hi-C matrices with 500 kb bins. Second, the Hi-C contacts are stratified by genomic distance, and a standard Pearson correlation is computed separately for each distance. Third, a novel statistic, the “stratum-adjusted correlation coefficient” (SCC), is computed as a weighted average of the distance-specific Pearson correlation, with weights derived using the Cochran-Mantel-Haenszel statistic. The SCC has a range of [1,−1] and is interpreted similarly to the standard correlation coefficient.

GenomeDISCO [21] performs a series of smoothing operations on the Hi-C matrices, and then calculates a pairwise similarity score separately at each smoothing level. The smoothing is performed using a network diffusion operation, where the nodes in the network are genomic loci and weighted edges are the contact counts in the Hi-C matrix. The diffusion operation effectively calculates, for each pair of loci *i* and *j*, the probability that a random walk will traverse the network from *i* to *j*. The smoothing level is controlled by raising the Hi-C matrix to the power *t*, where smaller values of *t* perform local smoothing and vice versa. The score for two matrices is the L1 distance (i.e., the sum of the absolute values in the difference matrix). These scores are summed across multiple values oft and then normalized to the range [−1,1], where 1 corresponds to identical matrices.

HiC-Spector [22] begins by computing, for each Hi-C matrix *M*, the corresponding Laplacian matrix *L = D−M*, where *D* is a diagonal matrix in which *D*_*ii*_ = *Σ*_*j*_ *M*_*ij*_. The matrix *L* is then normalized by the transformation *D*^−1/2^ *LD*^−1/2^, and its leading eigenvectors are found. The HiC-Spector score is defined as

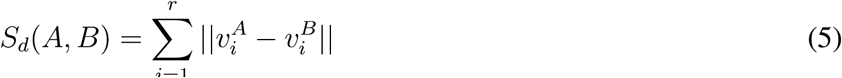

where 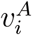 is the *i*th normalized leading eigenvector, and *r* is a user-specified parameter. The score is then linearly rescaled to the range [0,1]. In this work, the parameter *r* is set to 2.

We converted the three similarity measures to distances by treating each similarity as the cosine of the angle between two Hi-C matrices in some implicit high-dimensional space. This approach is justified because all of the eigenvalues of the similarity matrices are positive. Accordingly, the Euclidean distance between two Hi-C matrices *M*_*k*_ and *M*_*k′*_ can be computed as *d*(*M*_*k*_, *M*_*k′*_) = 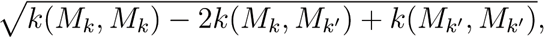 where *k*(*M*_*k*_, *M*_*k′*_) is a positive semidefinite similarity function.

We considered including several other possible measures, but ended up not using them. We did not use QuASAR [23] because the software was not working properly at the time we did our analysis. We also did use not the Pearson correlation coefficient (even though this measure has been used in some published studies [24-26]) because recent work shows that this approach does a markedly poor job of capturing relevant features of Hi-C data [19].

### 2.3 The similarity-based embedding approach

In this work, we focus on dimensionality reduction techniques, because projection to 2D provides an inuitive way to understand complex data sets. In general, most dimensionality reduction methods can be placed into one of two categories: *feature mapping* approaches versus *distance preserving* approaches. Feature mapping approaches directly find a mapping function from the original, high-dimensional space to a low dimensional space, subject to some optimization goal. A commonly used feature mapping technique is principal component analysis (PCA), which aims to find a linear transformation of a given set of features while preserving, as much as possible, the variance of the original data. In contrast, distance preserving approaches project the data points to a low dimensional space in which the original pairwise distances (or similarities) between pairs of data objects is preserved. Multidimensional scaling (MDS) [27] is a canonical example of a distance preserving approach. Although in principle a feature mapping technique can be applied directly to the Hi-C contact matrix by treating each matrix entry as a feature, such an approach is unlikely to accurately account for the complex features of Hi-C data. Such features include noise characteristics that are unique to Hi-C data, the tendency for pairs of loci that are close together along the genomic axis to come into contact frequently (the “genomic distance effect”), an topological features such as territories, domains and loops. To make good use of existing approaches like HiCRep, GenomeDISCO and HiCSpector that compute pairwise similarities between Hi-C matrices, we therefore chose MDS as our embedding approach.

The four sets of distances from Section 2.2 are provided as input to MDS, which is then used to project the data to two dimensions. We then order the cells by their projected angles. The projected angle *A*_*i*_ of cell *i* is the arc-tangent of its Euclidean coordinates *x*_*i*_ and *y*_*i*_, i.e., *A*_*i*_ = arctan(*x*_*i*_ − *x*_0_, *y*_*i*_ − *y*_0_), where (*x*_0_, *y*_0_) is the origin of the projection. We set *x*_0_ and *y*_0_ as the average of *x*_*i*_ and *y*_*i*_ across the cell population. In the end, we order the cells by their projected angles.

### 2.4 Assessing cell-cycle ordering

To evaluate the quality of a given method for ordering cells on the basis of Hi-C data, we need a way to quantify the agreement between an inferred ordering of cells and a given set of cell-cycle phase labels. To this end, we developed the “averaged circular ROC” that generalizes the standard receiver operating characteristic analysis to this multi-class, circular setting. The general idea is to treat each respective phase of the cell cycle in a one-vs-the-rest fashion, and to modify the usual ROC calculation to take into account the circular nature of the cell cycle.

More formally, the procedure proceeds as follows. Say that our inference method assigns a cell cycle angle to each of *n* cells (*θ*_1_, …, *θ*_*n*_), and that we are also provided with corresponding labels for each cell (*ℓ*_1_, …, *ℓ*_*n*_). For each cell, we know that 0 ≤ *θ*_*i*_ < 2*π* and that 1 ≤ *ℓ*_*i*_ < *m*, where *m* is the number of labeled phases of the cell cycle. To compute the circular ROC (CROC) score, we designate one label type as the positive class and all other label types as negative. This designation effectively transforms the multinomial labels into binary labels. For the positive class, we assume that the angles follow a von Mises distribution (also known as a circular normal distribution) and identify the mean angle *θ*\* as the maximum likelihood estimate of its mean parameter. Subsequently, we compute the minimal absolute difference between each cell’s assigned cell cycle angle and the mean angle: 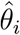 = min(|*θ*_*i*_ − *θ*\*|, |*θ*_*i*_ − 2*π* − *θ*\*|). The ROC calculation is then performed as usual, but using the binary labels and sorting cells according to 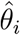. We then repeat this CROC calculation for each label type. The final score is the average of the area under curve of the CROC calculation (ACROC) across all *m* labels.

## 3 Results

### 3.1 Investigation of four potential distance measures

Initially, we evaluated the utility of the four distance measures (CDP-JSD, HiCRep, GenomeDISCO, HiC-Spector) based on their ability to project cells with known cell cycle phases. For this analysis, we selected 120 cells at random from the cell-cycle data set [14], including 30 cells from each of the four cell cycle phases (G1, early S, mid S and late S/G2). Applying our MDS embedding approach, we observe that HiCRep yields a projection that is markedly (albeit qualitatively) better than the other three: the resulting plot exhibits a clear circular pattern with the cells from the four cell-cycle phases placed in the correct order, G1 → early-S → mid-S → late-S/G2 → G1 (Figure 1). The MDS projection from GenomeDISCO also show some separation among the four cell-cycle phases, but no circular pattern is observed. In the projections produced by HiC-Spector and CDP-JSD, cells from different phases mix together.

**Figure 1:**
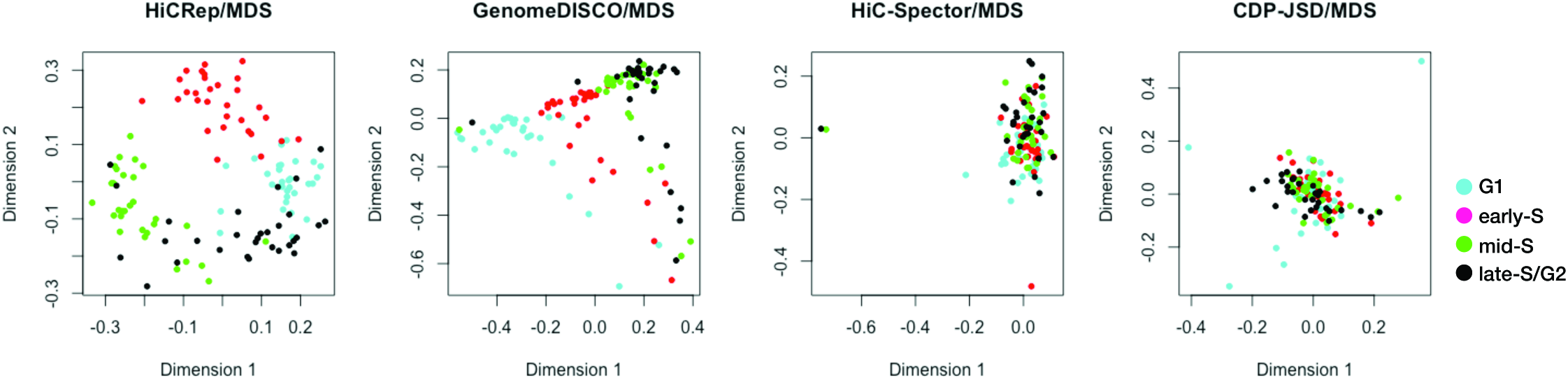
MDS projections of 120 cells from the four cell-cycle phases when the distance measure is calculated using HiCRep, GenomeDISCO, HiC-Spector and CDP-JSD.

Next, we developed a method (described in Sections 2.3–2.4) to assign cell cycle angles to each cell and then to quantitatively compare the inferred angles to the true phases derived from the orthogonal FACS and DNA content analysis. This evaluation yields a receiver operating characteristic curve for each phase of the cell cycle, along with an averaged score, the averaged circular ROC (ACROC). The quantitative results agree with the qualitative analysis: the ACROC achieved by HiCRep (0.940) is much greater than the scores achieved by the other three methods (0.816 for GenomeDISCO, 0.651 for HiC-Spector and 0.589 for CDP-JSD).

Based on this analysis, we selected HiCRep as our primary distance measure. In the rest of the manuscript, we further explore its utility in making sense of scHi-C data.

### 3.2 Comparison to a previously described cell phasing method

We next compare the cell-cycle phasing produced by our HiCRep/MDS projection with the phasing reported in Figure 2b of Nagano et al. [14]. For this analysis, we exclude the 120 cells used in previous experiment, focusing on the remaining 1051 cells from the cell-cycle data set. This cell-cycle phasing is derived from two different data sources: the contact distance profiles from the scHi-C data, and replication timing information from mouse Repli-chip data [28]. For each cell, Nagano et al. calculate a “replication score” based on the relative coverage of early-replicating regions, as determined from the Repli-chip data. Specifically, they classify each cell into one of five stages (post-M, G1, early-S/mid-S, mid-S/G2, and pre-M) by thresholding the proportion of near contacts (*p*near) and the proportion of mitotic contacts (*p*_mitotic_). They further order the cells within each of the five stages separately. For ordering cells in pre-M and post-M, they use *p*_mitotic_ only. For ordering cells in G1, they use *p*near and a score based on the mean contact distance among the contacts longer than 4.36 million basepairs. For ordering cells in early-S/mid-S and mid-S/G2, they use pnear and the replication score. As a baseline, we also include in our comparison the cell-cycle ordering produced by the aforementioned CDP-JSD/MDS approach, which represents the contribution from the contact distance profile only.

**Figure 2:**
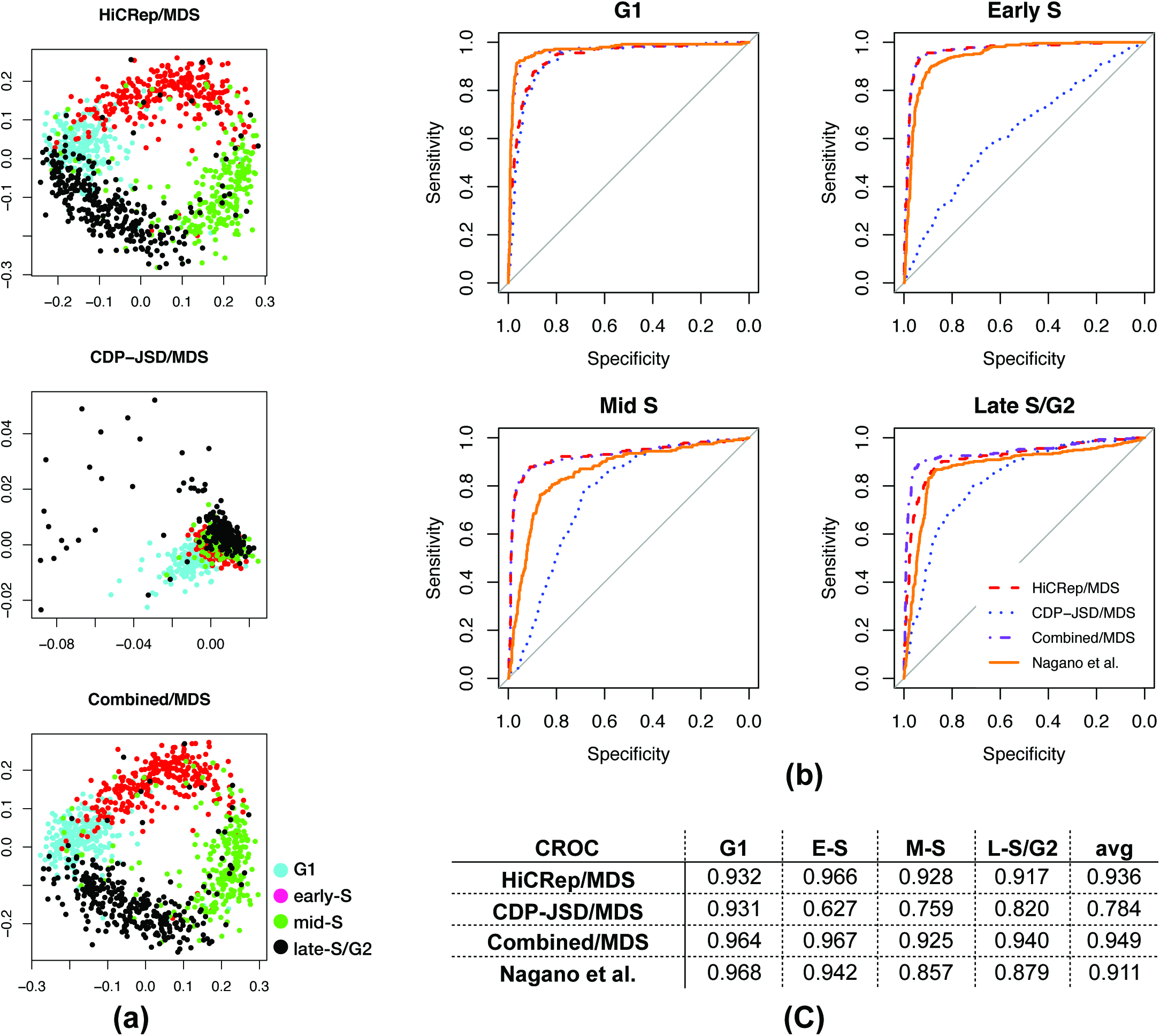
Comparison relative to the cell-cycle order from Nagano et al. (a) Projection of 1051 cells via HiCRep/MDS, CDP-JSD/MDS, and combined/MDS. (b) CROC curves from HiCRep/MDS, CDP-JSD/MDS, combined/MDS and Nagano et al. for the four cell-cycle phases. (c) Area under CROC curves from four approaches in the four cell-cycle phases, and the average CROC over the four cell-cycle phases.

This second set of projections confirms the utility of the HiCRep representation and suggests that the Nagano et al. procedure also produces accurate phasing (Figure 2A). Visually, the projection of the 1051 cells by Hi-CRep/MDS again yields a clear circular pattern with the cells from the four cell-cycle phases placed in the correct order. In contrast, the CDP-JSD representation yields a non-circular cluster. The ACROC score from HiCRep/MDS is 0.936, which outperforms the Nagano et al. cell-cycle order reported by 0.025 (Figure 2C). Investigating the CROC for the four cell-cycle phases, we observe that HiCRep/MDS outperforms the Nagano et al. method in the early-S, mid-S and late-S/G2 phases (0.966 vs 0.942, 0.928 vs 0.857, and 0.917 vs 0.879, respectively). HiCRep/MDS only underperforms the Nagano et al. method for the G1 phase (0.932 vs 0.968). The performance of the CDP-JSD/MDS method, which only uses the contact distance profile to evaluate distance, further confirms this result because it yields a high CROC for G1 (0.931) and low CROCs for the other three phases (0.627, 0.759 and 0.820 for early-S, mid-S and late-S/G2 phases, respectively). Thus, it seems that the contact distance profile information is particularly useful in identifying cells in G1, and that replication timing information is helpful in fully ordering cells according to their cell cycle phases.

Motivated by the performance of CDP-JSD/MDS in G1, we hypothesized that adding contact distance information to HiCRep may lead to improved performance. To test this hypothesis, we summed the HiCRep distance and the CDP distance and then applied MDS to the summed distances (termed *combined/MDS*). This combined/MDS approach further improves HiCRep/MDS for phasing cells in G1 phase and late-S/G2 phase by increasing the ACROC from 0.932 to 0.964 and from 0.917 to 0.940, respectively. The performance for early-S and mid-S does not improve, suggesting that contact distance profile information primarily helps to separate cells in G1 versus G2.

### 3.3 The method works even with very few contacts per cell

In contrast to the single-cell Hi-C protocol proposed by Nagano et al., which involves physically isolating individual cells and then amplifying and sequencing their DNA, single-cell combinatorial indexed Hi-C (sciHi-C) [16] uses a series of bar codes applied to cells that are randomly distributed in 96-well plates. The sciHi-C assay produces data from a much larger number of cells but characterizes each cell with much lower numbers of read pairs per cell. Therefore, we next investigate the robustness of the HiCRep/MDS methodology to reduced coverage in the Hi-C contact maps. In the 1051 cells used in Section 3.2, the number of contacts ranges from 20k to 654k, with a median 273k. We select the 20% of the cells with the lowest contact count (median 87.7k). We then randomly downsample the data so that each cell contatins 10k, 5k, 2k, or 1k contacts.

Projecting this series of five increasingly sparse data sets shows the separation of the four cell-cycle phases gradually disappearing as the number of contacts in the Hi-C contact maps decreases (Figure 3). Qualitatively, the initial set with median count 87.7k produces a circular pattern that resembles that produced by the full set in Figure 2a, but then the circular pattern decreases until it disappears entirely when the number of contacts drops to 2k or lower. This trend is captured quantitatively by the ACROC analysis (Figure 3f). All of four cell-cycle phases exhibit a similar degradation, although early-S is somewhat more robust to lower coverage. This observation suggests that the DNA structure changes occurring during early-S phase can be relatively easily captured by HiCRep. Overall, we conclude from this analysis that the HiCRep/MDS approach is quite robust and can provide a satisfactory projection as long as the number of contacts is above 5k.

**Figure 3:**
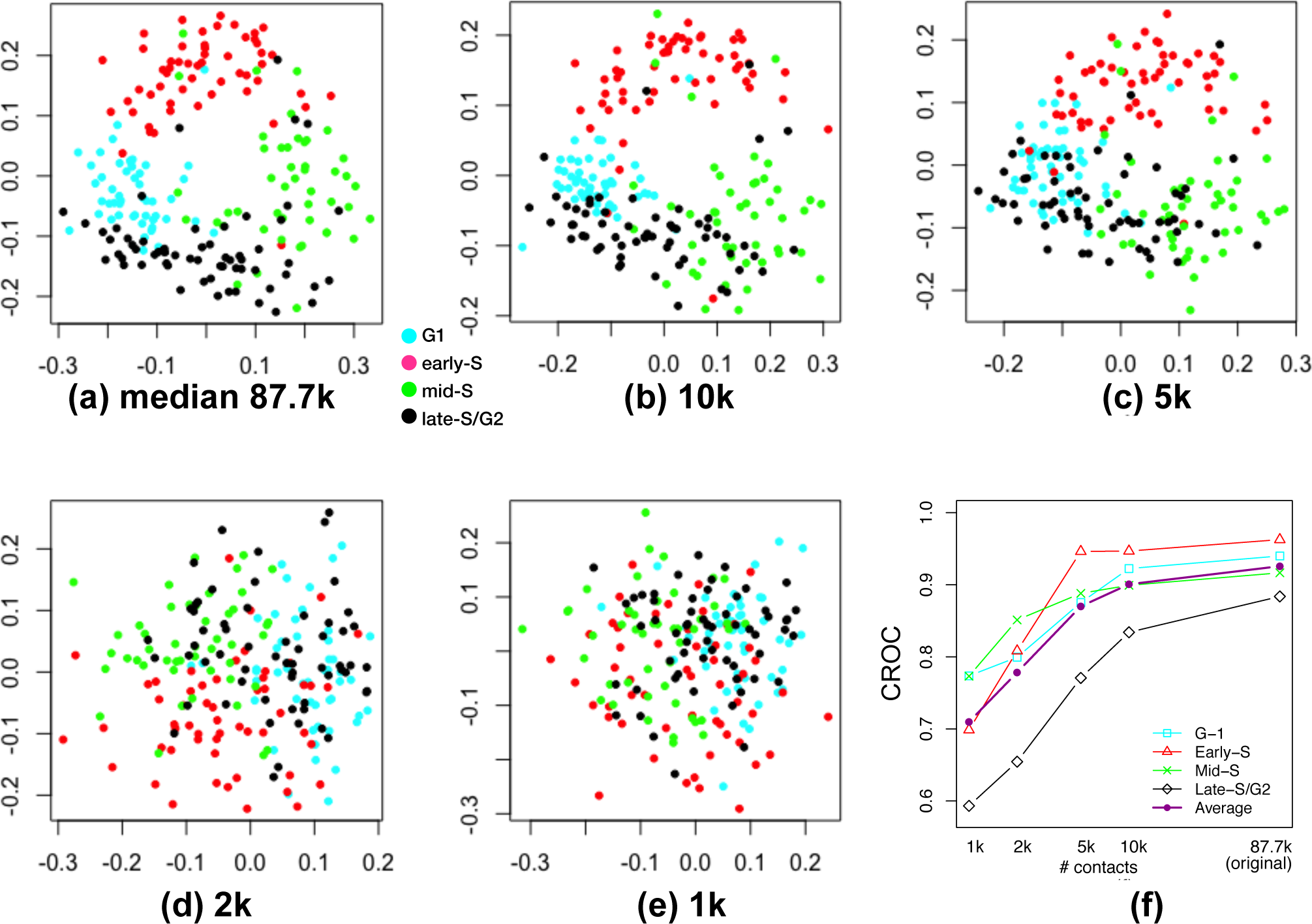
Downsampling analysis demonstrates that HiCRep/MDS projection is robust to decreasing coverage. (a) Projection of the low contact number set (bottom 20% with a median contact count 87.7k). (b-e) Projection of the same set of cells when they are downsampled to 10k, 5k, 2k and 1k, respectively. (f) The area under the circular ROC curves from different sets of downsampled cells for different cell-cycle phases (G1, early S, mid-S, and late-S/G2) and the average over the four cell-cycle phases.

### 3.4 Joint projection with high coverage cells improves phasing of low coverage cells

In practice, single-cell Hi-C protocols yield cells with a wide variety of sequencing depths. Accordingly, we want to know whether we can improve our ability to embed cells with low coverage by embedding them jointly along with other, higher coverage cells. We therefore projected the remaining 80% of the cells, along with the low-coverage cells downsampled to 1k, in a single projection (Figure 4a). Intriguingly, the downsampled cells form a concentric ring inside the ring formed by the original data set. Zooming in on the downsampled cells (Figure 4b), we can see that segregation of cells by cell-cycle phase is maintained in this “inner ring.” ACROC analysis (Figure 4c) confirms that including the higher coverage cells “rescues” the cell cycle phasing: when projected alone, the 1k cells achieve an ACROC of 0.710, whereas the same cells achieve an ACROC of 0.883 when projected jointly with the remaining 80% of the cells. To further test this idea, we split the remaining 80% of the cells into two equal sized groups (“mid” coverage and “high” coverage). ACROC analysis (Figure 4c) shows that including either one of these groups of cells along with the 1k set successfully rescues the ACROC of the 1k set. These results suggest that our proposed embedding methodology can be applied to scHi-C data from cells with varying sequence coverage, and that the presence of the higher coverage cells will help to properly embed the lower coverage cells.

**Figure 4:**
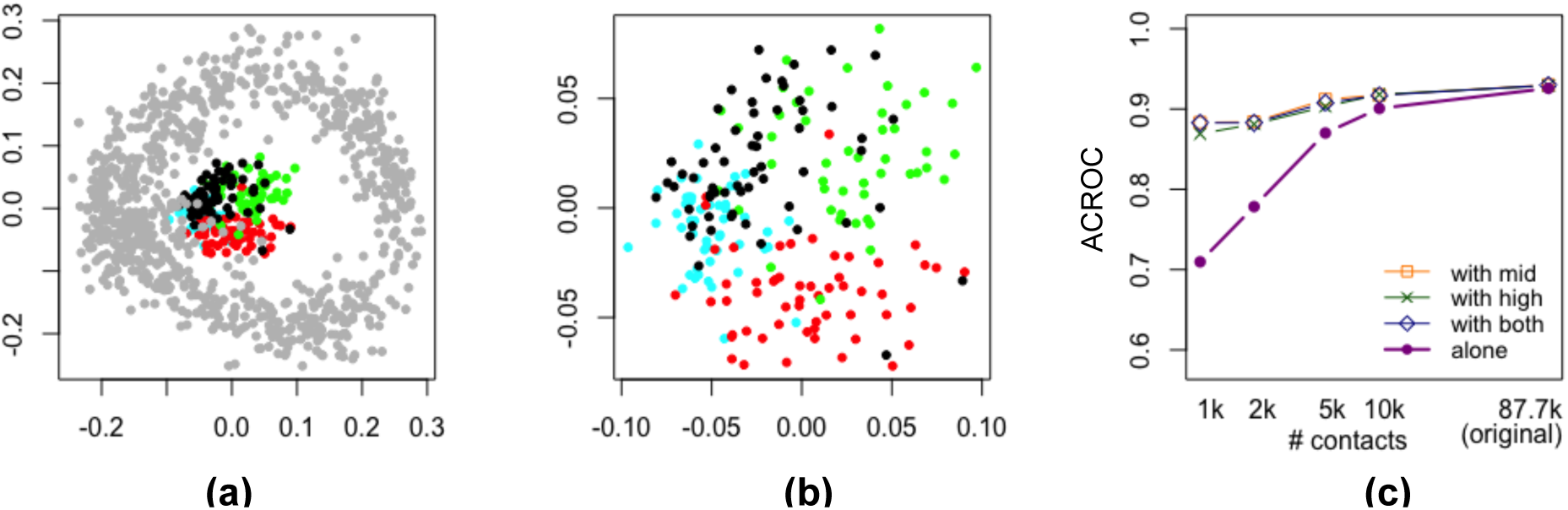
Joint projection of low quality cells with high quality cells. (a) Projecon of the cells which are downsampled o Ik (colored by their cell-cycle phase) together with the original high coverage cells (in gray). (b) Zoom-in view of the same projection of downsampled cells in (a), (c The area under the circular ROC curves from different sets of downsampled cells. Projections are performed on the cells by temselves, and in conjunction with the set of “mid” coverage cells, “high” coverage cells, or both.

### 3.5 Cell-cycle phased scHi-C data is indicative of replication timing

To further validate the quality of the cell-cycle phasing information inferred by the HiCRep/MDS approach, we investigate the relationship between a cell’s assigned cell-cycle phase and sequencing depth in regions of the genome with known replication times. The motivation is that as the cell enters the early S phase when DNA replication starts, the coverage of early replication regions from Hi-C data should increase, compared with the coverage of late replication regions. Conversely, when the cell reaches the late S phase when DNA replication is about to end, the coverage of the late replication regions will gradually catch up.

To test our hypothesis, we select two previously identified early replication regions (chr1: 36.5M-37M and chr1: 72.5M-73M) and two late replication regions (chr1: 20M-20.5M and chr1: 30M-30.5M) according to the *mus Musculus* 129ES-D3 Repli-chip (replicate 1) dataset [28]. We plot each cell’s coverage of this region versus the cell’s phase angle, as inferred by HiCRep/MDS. The early replication regions show a clear increasing-decreasing trend during S phase, and the coverage of the late replication regions shows the opposite decreasing-increasing trend (Figure 5). These results suggest that the inferred phasing agrees with prior knowledge about DNA replication timing.

**Figure 5:**
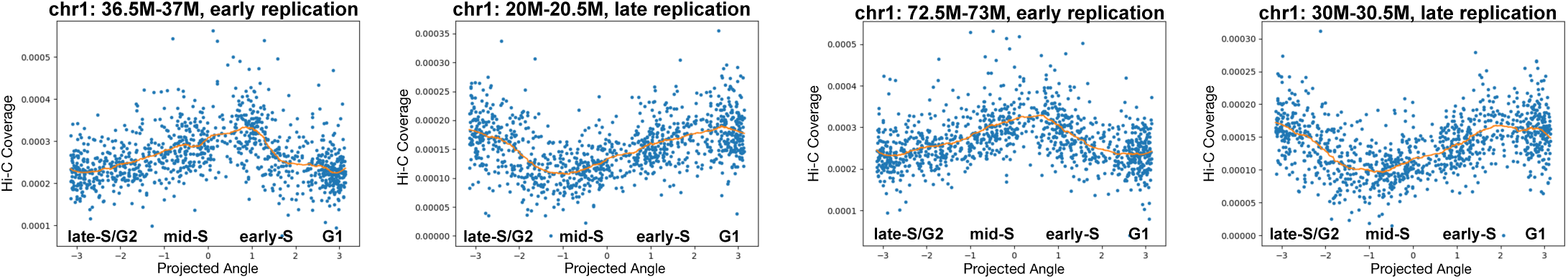
Inferred cell cycle ordering agrees with replication timing. Each panel plots the normalized Hi-C coverage (y-axis) of a specified genomic region as a function of the inferred cell cycle phase (x-axis). Two early replication regions and two late replication regions are depicted.

### 3.6 The method facilitates analysis across different cell types and experiment conditions

In preparation for an increasing number of publicly available scHi-C datasets, we want to investigate whether our embedding approach allows us to analyze multiple datasets together. For this purpose, we use two sets of scHi-C contact maps, the cell-cycle data set derived from mouse embryonic stem cells [14] and a data set derived from mouse cells during oocyte-zygote transition [15]. Unlike the cell-cycle data set, which has independently derived labeling of cell-cycle phase, the oocyte-zygote data set has no cell cycle labels; however, for this data set only zygotes in the G1 phase were scHi-C sequenced.

Projecting the two data sets jointly, we observe that the circular pattern from the mouse ESCs remains, and the four cell-cycle phases are clearly separated (Figure 6a). Thus, the cell cycle separation is robust even to mixing of these two different datasets generated from different scHi-C protocols. Also, the oocyte-zygote cells are projected adjacent to mESCs in G1 phase, with the oocytes and zygotes separated from one another. The male zygotes and female zygotes are also separated in the projection, whereas immature oocytes (NSN) and mature oocytes (SN) overlap to some extent. The proximity of the zygotes to G1 agrees with the reported cell cycle phase for these cells. While we do not know the cell cycle phase of the oocytes, the projection suggests that the DNA structure of oocytes is similar to mESCs in G1 or on the G1/early-S boundary.

**Figure 6:**
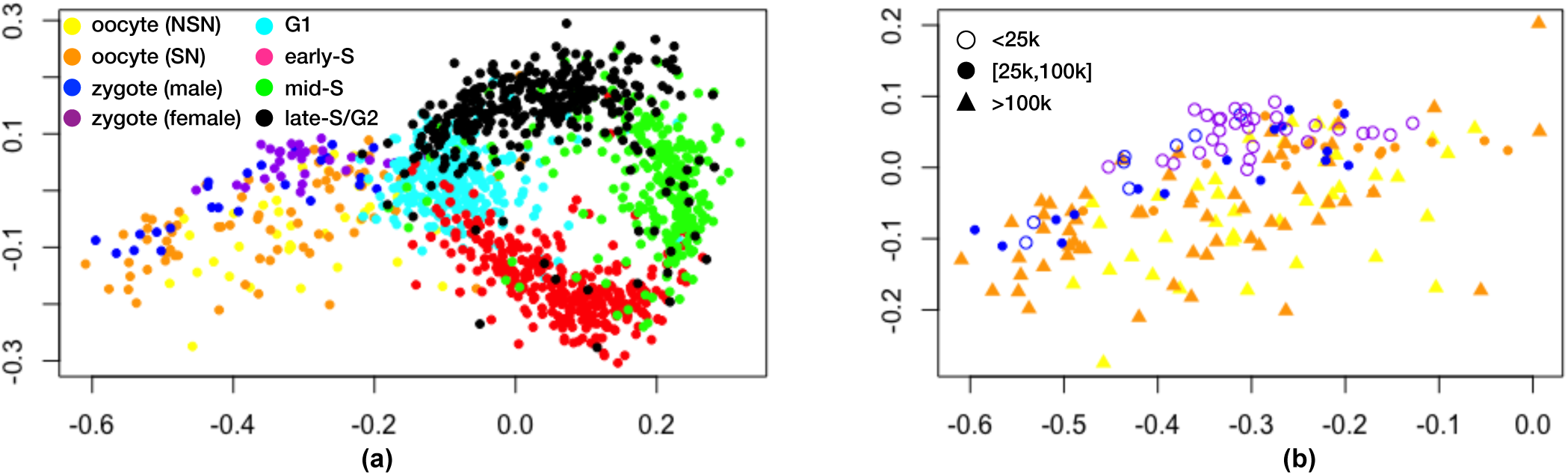
Joint HiCRep/MDS projection of cell-cycle and oocyte-zygote scHi-C data. (a) Joint projection, with cells from different stages colored differently. (b) The same projection, showing only the oocyte-zygote cells, with cells of different contact number levels labeled differently.

We also observe a large spread in the coordinates of the projected oocyte-zygote cells. This variability may be due to intrinsic variation in the 3D chromatin architecture of these cells, or it may be an artifact driven by the variation in sequencing depth in these cells. Indeed, the “double ring” in Figure 4a suggests that cell coverage may drive more deeply sequenced cells to be further from the center of the cell cycle ring. To investigate whether a similar effect is occuring in the oocyte-zygote projection, we re-plot the projected oocyte-zygote cells, labeling the points according to their sequencing depth. The result (Figure 6b) shows no obvious pattern with respect to contact number, suggesting that some other feature—possibly inherent stochastic variability in 3D structure— drives the observed variation in placement of the oocyte-zygote cells.

## 4 Discussion

We have demonstrated that HiCRep, used in conjunction with MDS, provides a powerful way to embed scHi-C data into a low-dimensional space that successfully captures biologically meaningful variation in 3D chromatin structure. In particular, the ability of this straightforward method to successfully separate cells by cell-cycle phase even from very low numbers of sequencing reads bodes well for the viability of sciHi-C, which yields many cells but lower average sequencing depth than the original scHi-C protocol. Ongoing efforts by large-scale projects such as the 4D Nucleome Network [29] promise to produce diverse scHi-C data sets. Our results suggest that these data can potentially be analyzed jointly using embedding methods such as the one proposed here.

Clearly, the current work is only a first step toward more sophisticated scHi-C analysis methods. For example, alternative embedding procedures, including t-SNE [30] and kernel PCA [31], may yield better embeddings. Particularly exciting is the potential to use kernel methods to jointly analyze multiple single-cell data modalities derived from the same population of cells. However, exploration of these avenues will require not only more scHi-C data, but also additional data sets that include orthogonal experimental characterization of the same single cells. Such data sets will enable supervised learning and also permit principled evaluation of competing methods.

